# Multiplexing bacteriocin synthesis to kill and prevent antimicrobial resistance

**DOI:** 10.1101/2024.09.06.611659

**Authors:** Alex Quintero-Yanes, Kenny Petit, Hector Rodriguez-Villalobos, Hanne Vande Capelle, Joleen Masschelein, Juan Borrero, Philippe Gabant

## Abstract

Antibiotic resistance represents an emergency for global public health. This calls for using alternative drugs and developing innovative therapies based on a clear understanding of their mechanisms of action and resistance in bacteria. Bacteriocins represent a unique class of natural molecules selectively eliminating bacteria. These secreted proteins exhibit a narrower spectrum of activity compared to conventional broad-spectrum antimicrobials by interacting with specific protein and lipid receptors on bacterial cell envelopes. Despite their diverse molecular structures, the commonality of being genetically encoded makes bacteriocins amenable to synthetic biology design. In using cell-free gene expression (CFE) and continuous-exchange CFE (CECFE), we produced controlled combinations (cocktails) of bacteriocins in single synthesis reactions for the first time. A first set of bacteriocin cocktails comprising both linear and circular proteins allowed the targeting of different bacterial species. Other cocktails were designed to target one bacterial species and considering bacteriocins pathways to cross the cell-envelope. Such combinations demonstrated efficient bacterial eradication and prevention of resistance. We illustrate the effectiveness of these bacteriocin mixtures in eradicating various human pathogenic-multiresistant—isolates. Finally, we highlight their potential as targeted and versatile tools in antimicrobial therapy by testing a combination of bacteriocins for treatment *in vivo* in the animal model *Galleria mellonella*.

## Introduction

The escalating issue of antibiotic abuse has precipitated a global crisis in antimicrobial resistance (AMR). The indiscriminate use and over-prescription of broad-range antibiotics have led to bacteria becoming resistant to conventional treatments, threatening the efficacy of our primary tools against bacterial infections. In the face of this crisis, it is imperative to develop innovative strategies based on alternative antimicrobial compounds.

Bacteriocins, a class of naturally occurring antimicrobial peptides (AMP) produced by bacteria, offer a distinctive solution in the fight against bacterial infections [1]. Bacteriocins can display both a broad- and narrow-spectrum of killing activity. Nonetheless, compared to antibiotics, many bacteriocins can target bacteria in a more specific manner. Therefore, it is expected that therapies based on bacteriocins targeting specifically pathogenic bacteria can help to prevent significant alterations in the microbiota (dysbiosis) [2] [3] [4] [5]. Indeed, targeting single bacterial species in communities provides competitive advantages for commensal bacteria to outgrow and displace pathogens if bacteriocin resistance cannot develop [2].

Synthetic biology emerges as an ally in maximizing the therapeutic potential of bacteriocins. Leveraging advanced techniques such as cell-free gene expression (CFE) and probiotic engineering, synthetic biology eases the production of antimicrobial peptides. Previously, we introduced the PARAGEN collection, a curated repository comprising bacterial-sourced DNA sequences standardized in silico for CFE, tailored to produce both linear and circular bacteriocins (**Figure 1**) [6] [7]. This collection is valuable for rapid screening, as it allows for the swift production (2–3 hours) and effective activity assessment of bacteriocins on bacterial lawns within a day (6–12 hours) (**Figure 1**). CFE utilizing bacteriocin genes from the PARAGEN collection has also been instrumental in elucidating the mechanisms of bacteriocin activity against human pathogenic bacteria [8] [9]. Moreover, *in vitro* expression has facilitated the synthesis of antimicrobial peptides (AMPs) from de novo coding sequences, designed using deep learning algorithms [10]. This approach yielded a substantial collection of short linear peptides, categorized based on their efficacy against bacteria and their low toxicity to human cells.

Probiotic engineering has emerged as a promising tool for harnessing bacteriocins to control bacterial communities [11]. Recently, non-pathogenic *E. coli* strains have been genetically modified to secrete bacteriocins that can target specific pathogenic bacteria, including ESKAPE pathogens [12] [13]. This innovative approach has demonstrated considerable potential, as the specificity of bacteriocins for the pathogenic species ensures the integrity of the engineered probiotics [12]. However, it is important to note that despite the concerted expression of bacteriocins, the pathogenic strains eventually developed resistance, outcompeting the probiotic strains over time [12].

The strategic use of antimicrobial combinations (also known as cocktails and mixtures) represents a significant advancement in overcoming challenges associated with bacterial infections. In the case of antibiotics cocktails, treatment potency can be augmented by synergy, rejuvenation, prevention of resistance and reduction of toxicity (reviewed in [14]). Similarly, antimicrobial cocktails composed of bacteriocins together with either antibiotics or phages have demonstrated the capacity to treat pathogenic bacteria ([15] and reviewed in [16] and [17]). Other cocktails composed solely of bacteriocins have proven to be effective in eliminating bacteria [18] [19]. It is noteworthy that combinations containing bacteriocins are commonly sourced with peptides synthesized either chemically or proteins produced *in vivo*, using ectopic gene expression and protein purification from bacterial pellets or culture supernatants. Compared to CFE, the chemical synthesis of large proteins can be problematic, while *in vivo* protein expression can be laborious and time consuming [20] [21].

A need for a rapid production and testing of bacteriocin combinations is required. In this research, we demonstrate that bacteriocin synthesis can be multiplexed in single CFE reactions to produce cocktails in a robust and fast manner. A first set of cocktails with bacteriocins of different sizes and structures allowed expanding the spectrum of activity to kill Gram + and - bacteria. In considering the bacteriocin pathways to cross the cell envelope, we synthesized other cocktails that kill bacteria effectively by synergy and prevented cross-resistance. Our work provides practical insights into developing targeted antimicrobial strategies that can be rapidly tested *in vitro* and *in vivo*, which is crucial given the growing challenges of antibiotic misuse and AMR pressure on our medical system.

**Figure 1.**
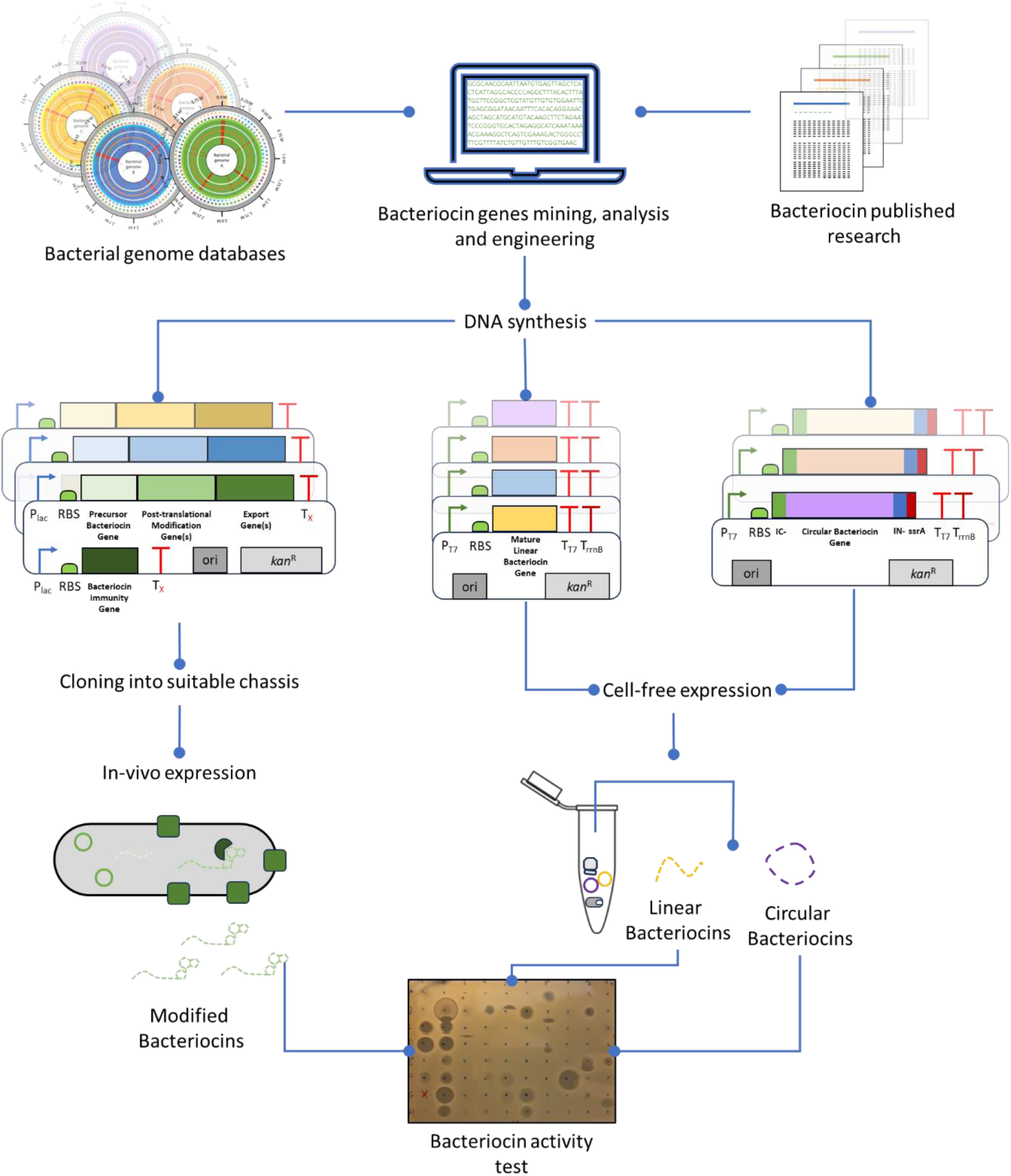
Bacteriocin *in vivo* and *in vitro* expression. Genes coding for bacteriocin peptides, post-translational modification, transport, and resistance are searched in bacterial genome databases and published research. Gene sequences are adapted for in-vivo heterologous expression and cell-free synthesis using synthetic biology approaches, such as standardized promoters, ribosomal binding sites (RBS), and terminators. All bacteriocin parts are synthesized de-novo and cloned into multi-copy vectors that can be transformed in *E. coli* for pDNA extraction. In the case of the cell-free expression, genes from the PARAGEN collection are under the control of the T7 RNA polymerase promoter, codon usage is adapted, and only the coding sequence for the mature peptide is synthesized, while for circular peptides bacteriocin genes are fused with inteins for circularization. Synthesis takes 12–16 hrs for *in-vivo* expression, and 2–3 hrs for *in vitro*. Thereafter, bacteriocins are tested on high-density bacterial cell lawns in their respective standard growth media and conditions.

## Results

### Cell-free co-expressed bacteriocins expand the spectrum of killing activity

In our endeavour to co-express bacteriocins within a single CFE reaction, we used recombinant transcription and translation proteins (PURE) alongside various vectors obtained from the PARAGEN collection. These vectors facilitated the expression of both linear and circular bacteriocins, including Enterocin L50A (EntL50A), Salmocin SalE1B (SalE1B), and circular bacteriocin Garvicin ML (GarML) (**Figure 2A**). EntL50A and GarML, derived naturally from *Enterococcus faecium* L50 and *Lactococcus garvieae* DCC43, respectively, are recognized for their targeting specificity towards Gram-positive bacteria such as *Pediococcus pentosaceus* and *Lactococcus lactis* IL1403 [22] [23]. Conversely, SalE1B, a colicin E1-like bacteriocin from *Salmonella*, demonstrates activity against closely related bacterial strains, including other *Salmonella* species and *E. coli* [24].

The efficacy of bacteriocins was assessed through agar diffusion assays, with spots applied on both Gram-negative and Gram-positive bacteria (*E. coli* DH10B or DH5α and *P. pentosaceus* CWBI B29, respectively). Individual bacteriocin synthesis reactions exhibited activity only against their expected bacterial targets (**Figure 2B**). Notably, when bacteriocins were synthesized collectively within the same reaction (CFE bacteriocin cocktail) we observed that both *E. coli* and *P. pentosaceus* strains were cleared (**Figure 2B**). Moreover, we noticed that adding a third DNA molecule coding for GFP did not compromise the killing activity towards both bacteria (**Figure 2B**). This demonstrates that bacteriocin CFE can be multiplexed to expand the spectrum of antimicrobial activity.

**Figure 2.**
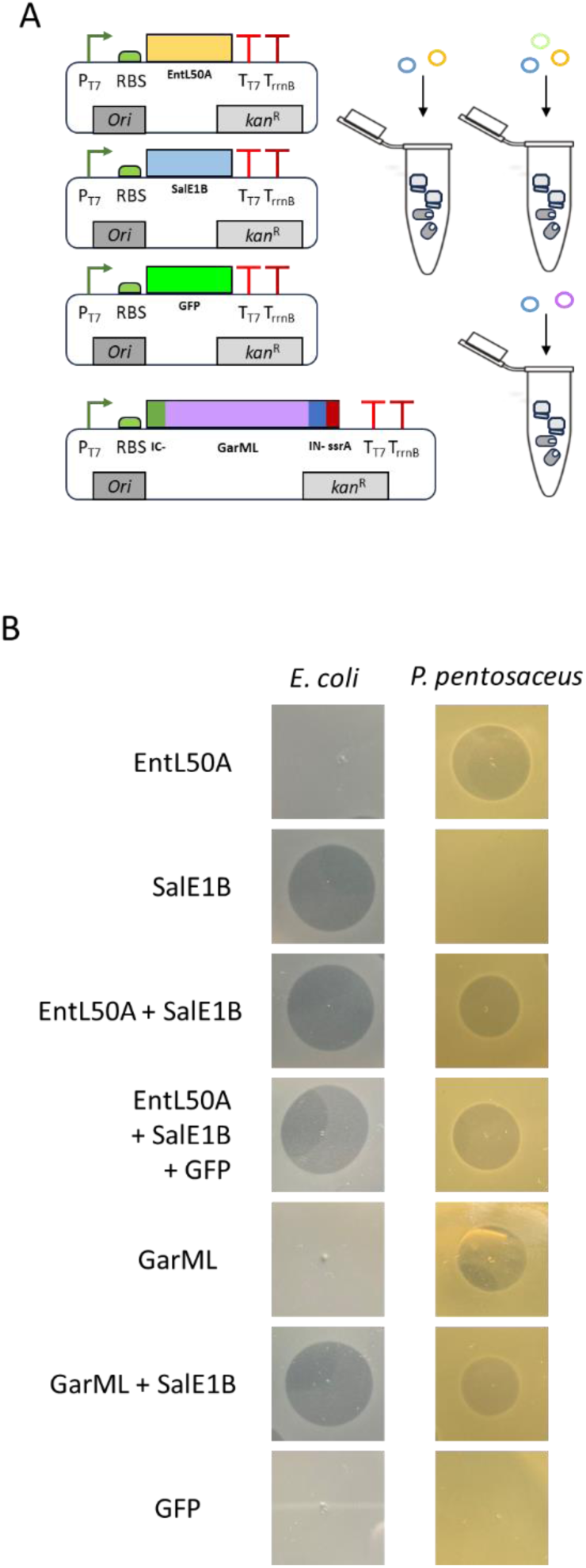
Co-expression of bacteriocins from PARAGEN in cell-free reactions. **A**. Vectors from the PARAGEN collection optimized for cell-free expression of either EntL50A, GarML, SalE1B or GFPmut3.1. **B**. Bacteriocin activity test on *E. coli* DH10B and *P. pentosaceus* CWBI-B29. Cocktails of EntL50A, SalE1B and GFP, and GarML with SalE1B were synthesized using PURE CFE.

### Rational bacteriocin cocktails increase killing efficacy

Considering that bacteriocin combinations synthesized in single CFE reactions retain activity, we tested whether two bacteriocins acting on one bacterial target could improve killing efficacy (i.e. in terms of synergy and AMR prevention). We chose to test such combinations against *E. coli* strains as the mechanisms underlying the activity of different bacteriocins have been extensively elucidated in this enterobacterial pathogen. Sensitivity to bacteriocins in *E. coli* hinges on the functionality of transport and structural proteins associated with the inner and outer membranes (**Figure 3A**). Notably, outer membrane porins exhibit lower specificity than inner membrane proteins (Reviewed by [25]). These porins rely on the Ton or Tol systems for their proper functioning, including interaction with bacteriocins. Consequently, single mutations in porins or Ton/Tol genes can confer pleiotropic resistance, leading to cross - resistance against multiple bacteriocins.

We investigated the CFE of combinations of various bacteriocins (microcins, colicins, and colicin-like) to assess their killing activity and prevention of antimicrobial resistance. First, we confirmed the activity of colicin M (ColM), SalE1B, microcin V (MccV) and microcin L (MccL) by complementing mutants from the KEIO collection with some of their cell-envelope receptor genes (**Figure 3A** and **Supplementary Figure 1A**). For instance, ectopic expression of *fhuA*, *cirA* and *sdaC* in mutants restored sensitivity to ColM, MccL and MccV, respectively. Although receptor and transporter genes for SalE1B have not been determined experimentally, based on its homology with Colicin E1, we predicted that SalE1B to be a BtuB-TolC-TolA-dependent bacteriocin (**Figure 3A**). Indeed, we could confirm that SalE1B depends on TolC for activity on *E. coli* (**Supplementary Figure 1A**).

**Figure 3.**
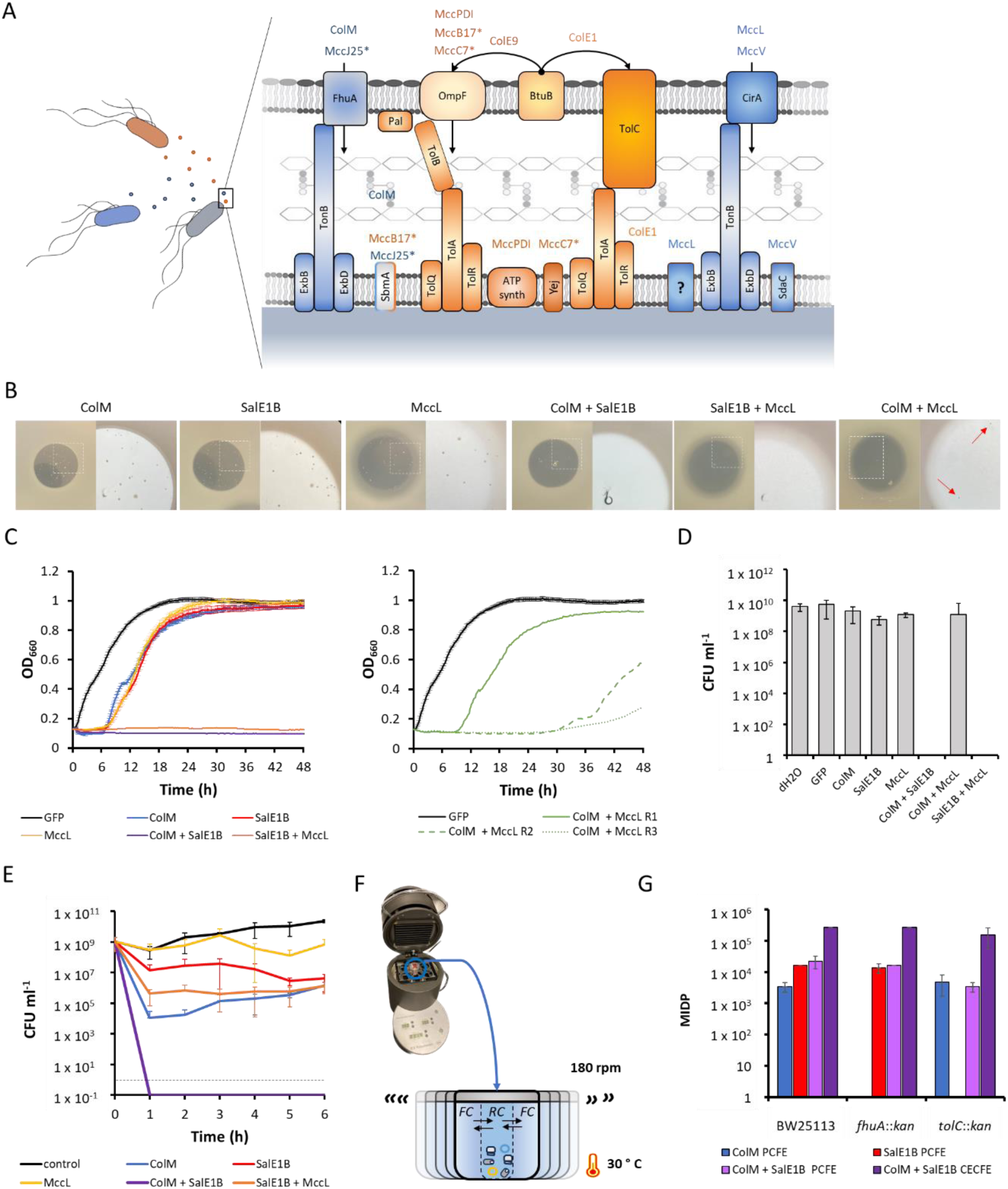
Rational designed and cell-free expressed bacteriocin cocktail. **A**. Bacteriocins pathways in *E. coli* cell-envelope. Tol and Ton-dependent bacteriocin pathways are highlighted in orange and blue, respectively, as well, producer cells and bacteriocins targeting Tol and Ton pathways. Different colicins and microcins are distinguished with the Col and Mcc prefix, respectively, while post-translationally modified bacteriocins are indicated with an asterisk (*). **B**. Activity of different bacteriocins against *E. coli* BW23113, halo of activity as seen as naked eye (left) and under a stereo microscope to detail colonies appearing in the halos (right). **C.** Growth of *E. coli* BW23113 in liquid cultures supplemented with single and co-expressed bacteriocins. **D.** Viability of cells in cultures (C) after 72h. **E**. CFU counts of *E. coli* BW23113 in liquid cultures supplemented with single and co-expressed bacteriocins diluted 1250 X. **F**. Diagram of synthesis of the ColM + SalE1B cocktail with CEFCE. The feeding chamber (*FC*) contains substrates and energy cofactors, while the reaction chamber (*RC*) holds the pDNA, and transcription and translation machinery. Reactions are shaken to improve diffusion of substrates into RC and of byproducts of protein synthesis out to *FC*. **G**. Minimum inhibitory dilution as seen on plates (MIDP) of cocktail preparations using CFE and CECFE reactions. Lines in C and E and bars in D and G represent average of three biological replicates, ± SD.

**Supplementary figure 1.**
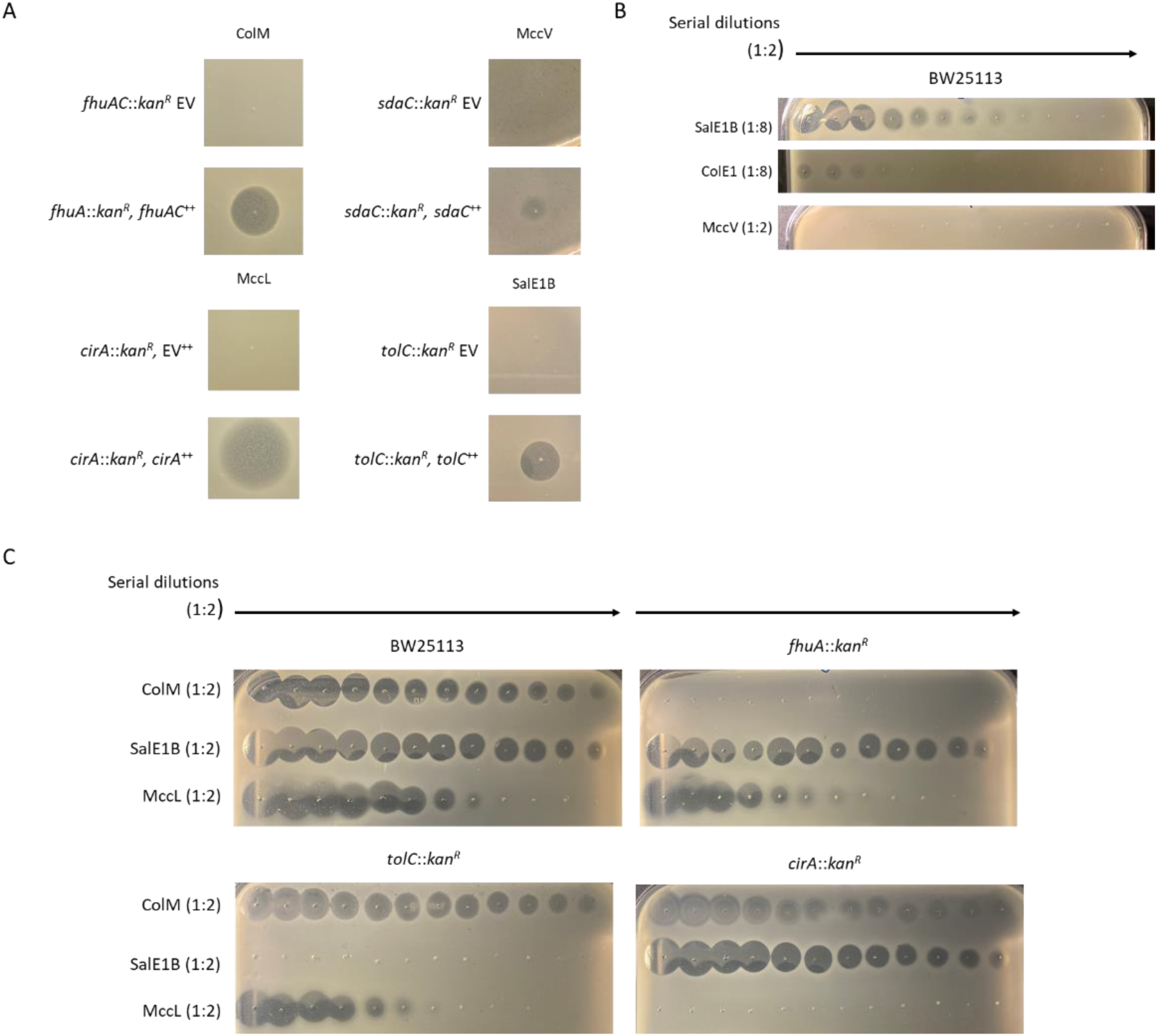
Activity of bacteriocins expressed with CFE. **A**. Activity test for Colicin M (ColM), Salmocin SalE1B (SalE1B) and Microcin L (MccL), Microcin V (MccV) on lawns of mutants for bacteriocin receptor genes from the KEIO collection carrying an empty vector (EV) and complemented with WT allele. The heterologous expression of genes was induced with 0.5 mM IPTG. **B**. Comparison of activity of SalE1B and ColE1 using serial dilutions (1:2). ColE1 and SalE1B were first diluted 1:8, while MccV was diluted 1:2. **C**. Activity of ColM, SalE1B and MccL using serial dilutions (1:2). All bacteriocins were first diluted 1:2.

We could also observe that SalE1B when synthesized with CFE, had a higher activity than ColE1 as seen in the activity of serially diluted bacteriocins (**Supplementary Figure 1B and C**). Other bacteriocins, such as ColM and MccL retained activity after serial dilutions, while MccV lost it (**Supplementary Figure 1B and C**). Consequently, we proceeded to prepare bacteriocin combinations using CFE with SalE1B, ColM, and MccL.

We monitored bacterial growth upon treatment with individually expressed bacteriocins. After overnight incubation, we observed the emergence of colonies within the halos of activity, accompanied by the resumption of growth in liquid cultures after 6 hours of arrest (**Figure 3B** and **C**). This phenomenon indicated the rapid development of resistance in our single bacteriocin assays in some cells within the colonies and cultures. Further characterization of the colonies within the SalE1B halo revealed gene mutations associated with the colicin E1 homologous pathway, such as *btuB*, *tolC*, and *tolA* (**Supplementary Figure 2**). Additionally, we complemented a mutant that exhibited complete insensitivity to SalE1B with *tolC* from the parental strain BW25113. Our findings support the notion that resistance stemming from mutations in receptor genes can readily emerge following a single bacteriocin treatment *in vitro*.

**Supplementary Figure 2.**
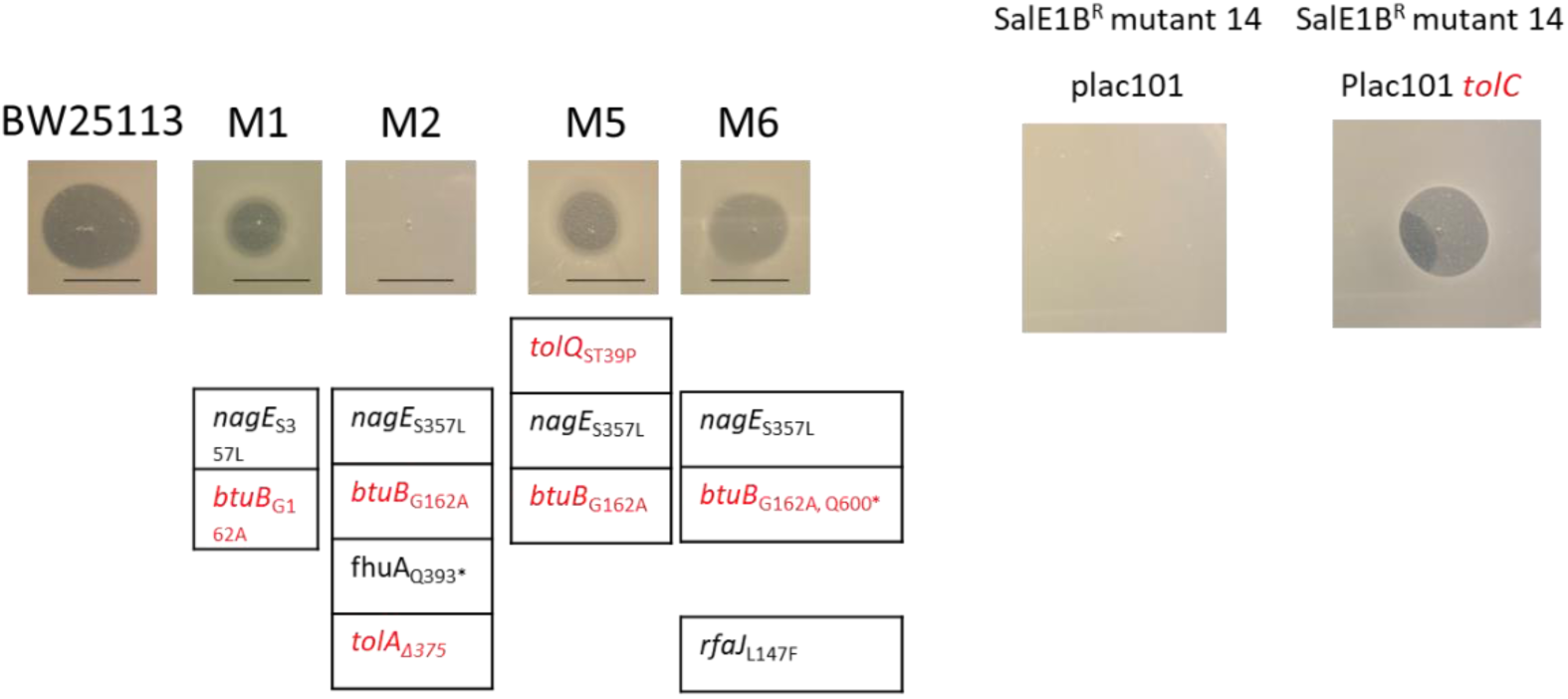
Mutations in expected cell-envelope genes are causing resistance. Colonies grown within the halo of SalE1B activity were isolated, tested against SalE1B and analysed using whole genome sequencing. One of the mutants showing complete insensitivity (mutant 14 with SNP *tolC_Q347*_*) was complemented with the *tolC* WT allele and sensitivity to SalE1B was restored. Scale bar in pictures: 1 cm.

Crucially, we noted that treatment with cocktails containing bacteriocins that utilize distinct pathways to enter the cell envelope (ColM + SalE1B and SalE1B + MccL) did not exhibit resistance (**Figure 3B** and **C**). Viability assays conducted on cells grown in liquid cultures further demonstrated that combinations ColM + SalE1B and SalE1B + MccL effectively eradicated the cells after 72 hours (**Figure 3D**). In contrast, when cultures were treated with ColM + MccL, which utilize the TonB-dependent pathway for activity, colonies appeared within the halo of activity, growth was observed in liquid culture, and viable cells were present (**Figure 3B-D**). This confirmed the importance of a rational design (considering cross-resistance) for bacteriocin cocktail preparations.

We diluted more than 1000 times the single and co-expressed bacteriocin solutions and tested against cultures of BW25113 to assess synergic effects (**Figure 3E**). Consistent with the activity on plates, MccL was not efficient in killing bacteria in such dilutions, while SalE1B and, especially, ColM, had better activities, compared to the cultures not treated with bacteriocins (control). It was observed that the co-expressed cocktails ColM + SalE1B and SalE1B + MccL had better killing activities than their respective individually synthesized bacteriocins, indicating that the bacteriocins co-expressed have a synergistic effect.

Recognizing that cell-free expressed ColM and SalE1B showed higher antibacterial activity, and that their combination effectively prevents resistance, we explored the ColM + SalE1B cocktail synthesis using a continuous exchange cell-free expression (CECFE) method (**Figure 3F**). CECFE facilitates the continuous flow of energy co-factors, ribonucleotides, and amino acids into the reaction compartment, efficiently removing by-products. First, we determined the yield of bacteriocin activity for single and co-expressed ColM and SalE1B with PURE CFE on plates with lawns of indicator strains (parental and keio mutants) (**Figure 3G**). This permitted confirming that the activity of the individual bacteriocins can be traced with specific indicator strains. More importantly, we could compare the activity of the cocktail produced with CECFE with the one with PURE CFE and determine that there is more than a 10-fold change in bacteriocin activity with the CECFE method. This demonstrates that the antimicrobial activity of bacteriocins cocktails can be scaled up.

Thus far, we have tested the activity of each bacteriocin in cocktails by using indicator bacterial strains displaying sensitivity to one component but not the other. To further characterize the ColM + SalE1B cocktail, we demultiplexed the bacteriocins using a direct activity test with samples fractionated in a Tris-Tricine-SDS page gel. This revealed that the bacteriocins in single and combined CFE preparations were in the predicted range of molecular weight, 29 and 58 KDa for ColM and SalE1B, respectively (**Figure 4A**). Furthermore, this confirmed that bacteriocins retained their linear active form and weight in the cocktail as observed in single reactions.

**Figure 4.**
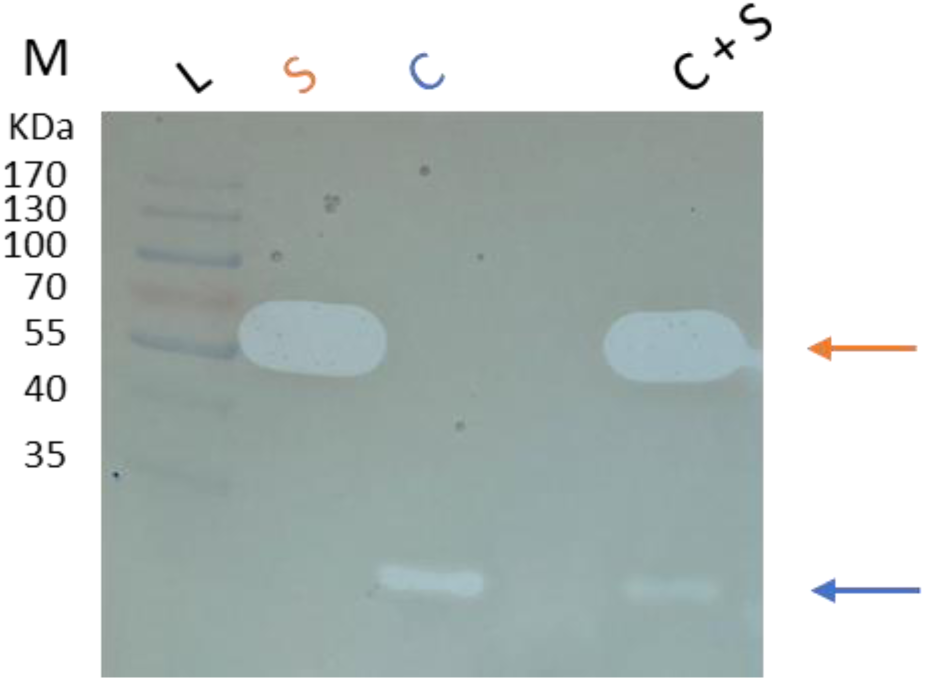
Separation of bacteriocins in cocktail. The activity of each bacteriocins in cocktail was distinguished following separation by molecular weight with Tris-Tricine SDS-page and test of activity on a top lawn of *E. coli* BW25113. L: ladder, S: SalE1B, C: ColM and C + S: ColM + SalE1B cocktail. The blue and green arrows indicate the predicted ColM and SalE1B bands, respectively.

### Bacteriocin cocktail to control infections with antibiotic resistant pathogenic strains in *Galleria mellonella*

We proceeded to test the bacteriocin cocktails to treat infections of clinical isolates *in vivo* in the animal model *Galleria mellonella*. First, we determined the toxicity of CFE cocktail preparations and observed that 10^2^ and 10^3^ dilutions did not kill and were less harmful to *G. mellonella* larvae than a more concentrated solution (**Figure 5A** and **B**).

Considering that a 10^2^-fold diluted ColM + SalE1B cocktail did not kill *G. mellonella* larvae and had minor secondary effects as seen in the health score assessment, we tested this cocktail dilution *in vitro* against lawns of clinical isolates of multiresistant pathogenic *E. coli*. These strains produce different AmpC and extended spectrum *β*-lactamases including CMY-2, SHV-11, CTX-M-14, CTX-M-15, and KPC, NDM, OXA-48 carbapenemases. We could observe that several strains were more, similar or less sensitive to ColM + SalE1B CFE preparations, compared to a lab strain DH10B, while others were completely insensitive to the cocktail (**Supplementary figure 3A**).

Thereafter, we assessed the pathogenicity of clinical isolates against *G. mellonella*. We observed that the selected strains were harmful to *G. mellonella*, leading to sickness and death within a day of infection (**Figure 5C** and **Supplementary figure 3B**). Notably, the cocktail proved to be an effective treatment for some selected isolates, including strains 141, 425, and 10276-2, as it helped maintain survival and health scores like those of the control population (larvae treated with saline solution). For the treatment against the other strains, the probability of survival and health score decreased, compared to the control population. However, these values were still significantly higher than those of larvae that did not receive treatment with the bacteriocin cocktail (**Figure 5C** and **Supplementary figure 3B**).

**Figure 5.**
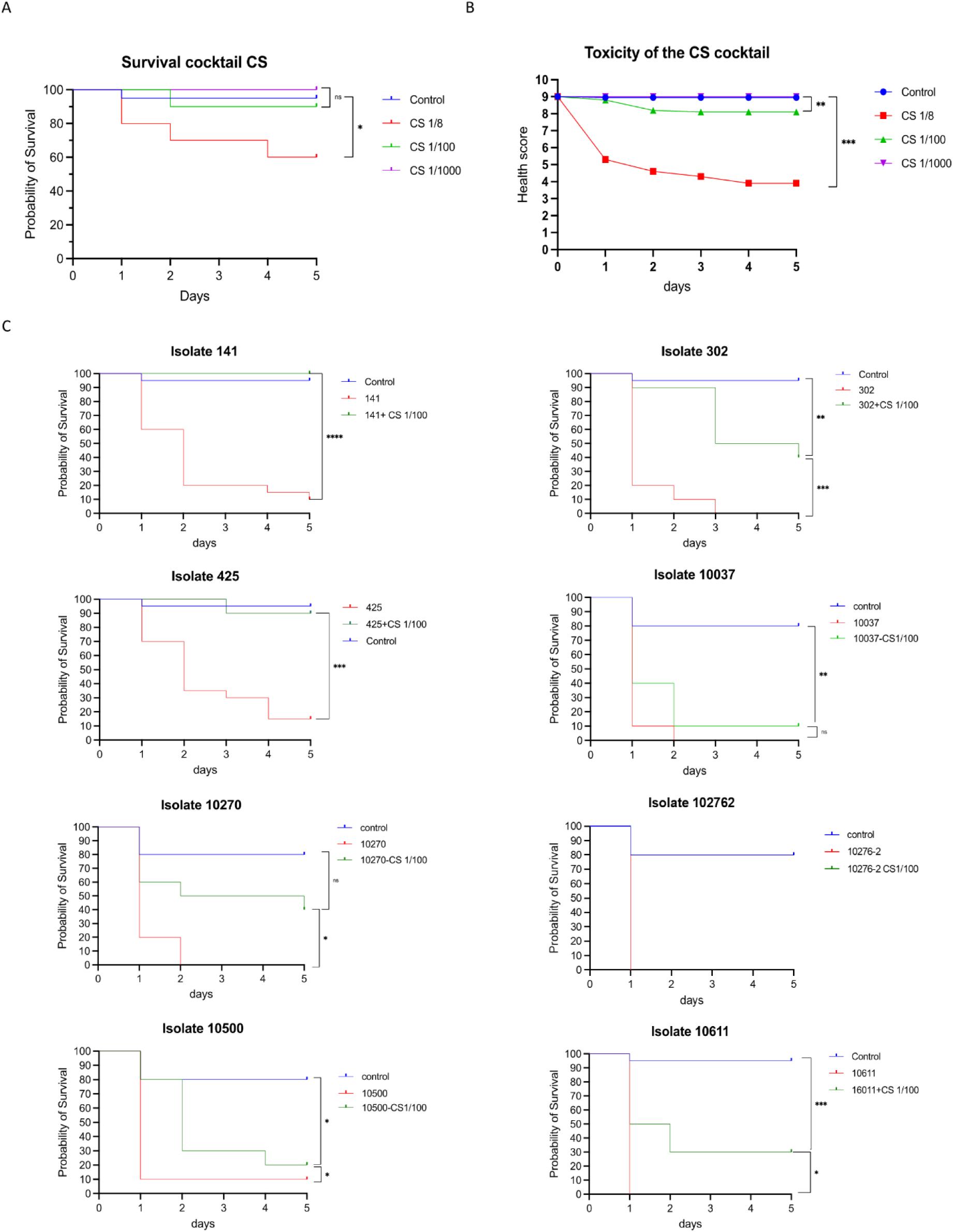
Diluted bacteriocin cocktail is not toxic and improve survival of *Galleria mellonella* after bacterial infection. **A-B**. Survival (A) and health (B) scores measurements on the animal model *G. mellonella* individuals after injection of different dilutions (1:8, 1:100 and 1:1000) of the ColM + SalE1B cocktail (C + S) produced by CECFE. **C**. Survival score measurements on *G. mellonella* after infection with antibiotic multi-resistant *E. coli* strains (141, 302,425, 10037, 10270, 10276-2, 10500, 10611). Bluelinein graph for treatment with isolate 10276-2 is superimposed on green one. n=10 per treatment, ns: not significantly different (*p* ≥ 0.05), * *p* < 0.05, ** *p* < 0.01 and *** *p* < 0.001.

**Supplementary Figure 3.**
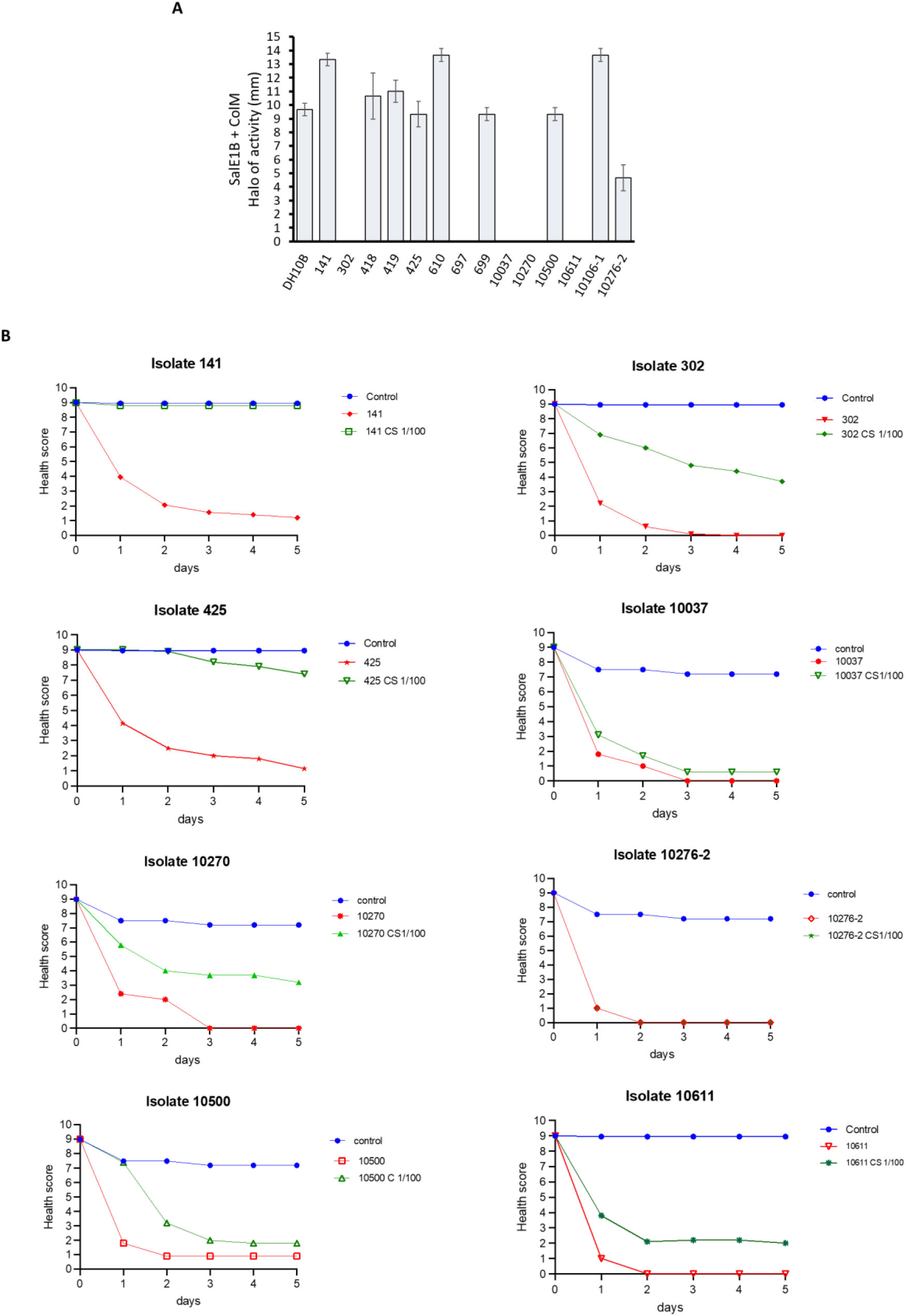
Activity *in vitro* and *in vivo* of ColM + SalE1B cocktail. **A.** Halo of cocktail activity on lawns of *E. coli* strains. Bars are representative of average value of three biological replicates, ± SD. **B**. Health scores measurements on the animal model *G. mellonella* individuals after injection of diluted (1:100) ColM + SalE1B cocktail (C + S) (from CECFE preparations) for treatment against antibiotic resistant clinical isolates. Blue line in graph for treatment with isolate 10276-2 is superimposed on green one. n=10 per treatment, ns: not significantly different (*p* ≥ 0.05), * *p* < 0.05, ** *p* < 0.01 and *** *p* < 0.001.

### Optimization of bacteriocin synthesis

Previously, we observed that cell-free expressed MccV showed poor activity compared to ColM, MccL and SalE1B (Supplementary Figure 1B). To improve MccV CFE, we re-designed gene parts from the PARAGEN collection, such as P_T7_ promoter, genes, and terminator sequences, using guidelines for template DNA design from PureFrex (GeneFrontier Corporation). First, we tested the optimization using *gfp* and observed a 10-fold change in fluorescence detection (**Supplementary Figure 4A**). Consequently, we re-engineered the original PARAGEN *mccV* device and tested the bacteriocin synthesis. Compared to the original PARAGEN devices, MccV from optimized DNA samples could be detected in samples diluted more than 200 times.

MccV is predicted to form disulfide bonds (DSB) in nature [26]. Therefore, we tested the synthesis of MccV using commercially available supplements to enhance DSB formation in CFE. Notably, the DSB enhanced CFE resulted in 2-fold change of MccV activity in the optimized device (**Supplementary Figure 4B**). Our results confirm that our PARAGEN collection can be re-engineered to achieve significantly higher bacteriocin activity, and that bacteriocin CFE conditions can be adapted for further improvement on activity.

**Supplementary Fig 4.**
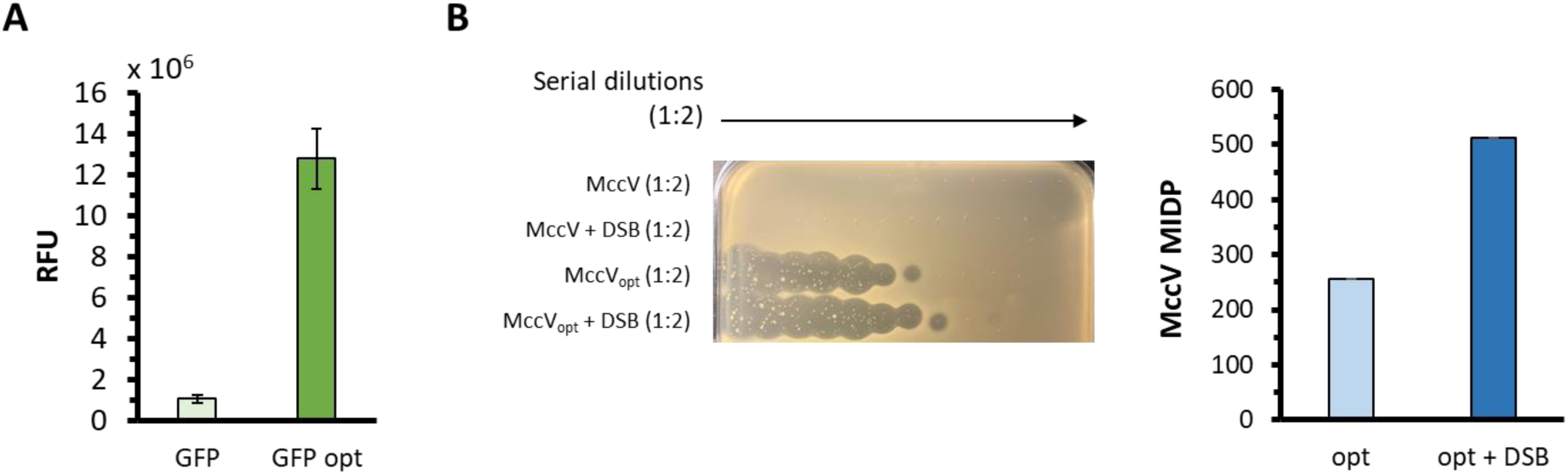
Optimization of PARAGEN sequences increase bacteriocin activity. A. CFE of GFP with (_opt_) and without optimization of DNA devices. **B**. Activity of serial diluted CFE bacteriocin samples with and without sequence optimized DNA devices and supplements for disulfide bond formation (DSB) against a top law of *E. coli* BW25113. MIDP of CFE MccV with optimized PARAGEN device without and with DSB. Bars are representative of the average value of three biological replicates, ± SD.

## Discussion

Here, we have shown that engineering and synthetic biology principles, such as standardization, optimization and multiplexing can lead to fast, robust, safe and customized combined therapies to treat pathogenic and antibiotic-resistant bacteria. Genetic devices from the PARAGEN collection ease the *in vitro* synthesis of bacteriocins with different sizes and structures. In the same conditions used to synthesize single bacteriocins, we efficiently co-expressed different proteins coded from different DNA molecules (multiplexing). In addition, we demultiplexed the signals (bacteriocins activity) in cocktails physically by using electrophoretic separation in non-denaturing conditions (Tris-Tricin-SDS-PAGE) and biologically by testing their activity against different KEIO collection strains sensitive to each bacteriocin specifically. Thus, we demonstrated that independent bacteriocin activity in cocktail preparations can be easily tested and validated.

More importantly, we have shown that treatments with bacteriocin combinations co-expressed in single CFE reactions are efficient in killing bacteria, expanding the spectrum of activity and preventing AMR. Also, we demonstrate that a CECFE for bacteriocin co-expression helps in scaling up cocktail synthesis and facilitates using diluted CFE bacteriocin solutions to control infections without several toxic effects in the animal model *G. mellonella*. On top of this, the Design-Build-Test-Learn cycle led us to re-engineer the PARAGEN collection and include posttranslational additives, such as a DSB enhancer blend, to increase bacteriocin production. Our work serves as a proof of concept to synthesize more complex bacteriocin combinations that could help to kill problematic bacteria.

In the case of the SalE1B + ColM and SalE1B + MccL cocktails here presented, we observed that the combinations worked better in killing activity and AMR prevention than single bacteriocins. Unlike treatments with single antibiotics, cocktails can lead to synergistic effects, rejuvenation of old antibiotics, reduction of resistance events and toxicity, and amplification of the spectrum of activity to tackle community-acquired infections [27] [28] [29]. It’s important to keep in mind that antimicrobial resistance (AMR) is on the rise, and the limited discovery and availability of new antibiotics hinders the development of alternative and efficient single and combined treatments. Therefore, alternative antimicrobials will soon become a source of therapy.

The in vivo bactericidal efficacy of a bacteriocin cocktail was demonstrated against highly resistant pathogenic *E. coli* strains harboring critical resistance genes, including CTX-M-15, OXA-48, KPC, and NDM. These findings create opportunities for the development of targeted bacteriocin cocktails as adjuvants in antibiotic therapies and facilitate the decolonization of specific resistant strains. Furthermore, they present promising new therapeutic options for infection control and the decontamination of medical devices contaminated by epidemic organisms. Therefore, further research is warranted to confirm the synergistic effects of these cocktails with antibiotics, disinfectants, and antibiofilm activities.

Co-expression of several gene devices and circuits using CFE has been described before [30] [31] [32] [33] [34]. Recently, the CFE of Salivaricin B, a bacteriocin with post-translational modifications (class I bacteriocin), was achieved using three different DNA molecules to co-express bacteriocin precursor, modification and maturation proteins [35]. Although our work here was restricted to the expression of bacteriocins with circular and linear structures out of single gene parts, it is likely that, future antimicrobial cocktails will be composed with CFE bacteriocins with more complex structures. It is even conceivable to think of CFE bacteriocin combinations with other antimicrobial proteins or entities co-expressed in single reactions. Indeed, the co-expression *in vitro* of enzymes for the synthesis of non-ribosomal antimicrobial peptides has been achieved, as well as the assembly of phages [36] [37].

## Methods

### Cell-free bacteriocin synthesis

*In vitro* bacteriocin synthesis was performed using bacteriocin genes in pDNA from the PARAGEN collection [6] [7] with PurExpress (New England BioLabs) and RTS 500 ProteoMaster *E. coli* HY (biotechrabbit) for CECFE. Synthesis conditions were followed according to each manufacturer’s recommendations. pDNAs for CFE of 2 or 3 proteins (bacteriocins + GFP) in single reactions were mixed in 1:1 or 1:1:1 ratio of concentrations, using the same concentration for single bacteriocin synthesis.

### Optimization of cell-free bacteriocin synthesis

Bacteriocin expression devices from the PARAGEN collection were optimized following template DNA design guidelines for CFE with PUREfrex (GeneFrontier Corporation). Our DNA templates were redesigned considering codon usage, AT content after the start codon, and removal of mRNA secondary structures and sequences likely causing frameshift during elongation. In addition, a *lac* operator sequence downstream of the T7 promoter was removed. The sequences for GFP and MccV expression prior and after optimization have been shared as supplementary files to provide detailed information.

### Bacterial growth conditions and strains

*E. coli* cells were grown at 37 °C for 6-16 h in either LB or Mueller Hinton medium, as specified. *Pediococcus pentosaceus* CWBI B29 cells were grown in MRS media for 18-24 h at 30 °C. *E. coli* strains with mutations for the different bacteriocin receptors were obtained from the Keio collection [38]. Growth assays in liquid cultures were done in 96 well-plates OD_600_ reads were taken every minute for 48 hrs using a SpectraMax i3X plate reader at 37 °C with linear shacking before each timepoint.

### Bacteriocin activity tests

The activity of bacteriocins was tested on plates and liquid cultures. On solid media, 2.5 µl taken either directly from the cell-free reactions or serial dilutions were spotted on plates with indicator strains. Plates were prepared by mixing fresh overnight cultures of either *E. coli* or *P. pentosaceus* normalized to 1.0 OD_600_ with 0.7 % agar in a 1:80 and 1:10 ratio, respectively, and pouring the mix on solid growth media (1.5 % agar). Bacteriocin spots on bacterial lawns were allowed to dry before incubation.

In liquid media, fresh seed cultures were diluted to 0.1 OD_600_ and the bacteriocin cell-free reactions were diluted 8 times (1:8). Then, 198 µl of the diluted culture was mixed with 2 µl of the diluted bacteriocins in 96-well plates, incubated and assessed for optical density as indicated above. CFU counts for viability test were done similarly, except that incubation was done using an orbital shaking incubator and samples were taken every hour.

The bacteriocin activity on *E. coli* clinical isolates was evaluated using Mueller Hinton agar plates (BD, USA) from pure cultures of each strain, in accordance with the EUCAST (European Committee on Antimicrobial Susceptibility Testing) guidelines for antimicrobial susceptibility testing by disc diffusion. A 2.5 µL spot of each bacteriocin was applied directly onto plates inoculated with a 0.5 McFarland suspension of a pure overnight culture strain. The diameter of the inhibition zones around each spot was measured after 24 hours of incubation at 35°C in an aerobic atmosphere.

### MID determination

MID is defined by the lowest dilution showing antimicrobial activity. To avoid any bias in determining the MID especially when the inhibition halos become faint, we developed a custom python script to automatically recognize and analyse these halos from plate images. The script identifies halos by detecting circular areas with a radius greater than 7 pixels and a pixel intensity value that is at least 5 units lower than the background pixel intensity. After testing and iterating with our image set, we found that setting the intensity difference parameter (x) to 5 provided the best balance between sensitivity and specificity.

### Fluorescence measurements

GFP fluorescence during cell free expression was performed using 25 µl reactions with PurExpress (New England BioLabs) in 384-well black microplates, with flat bottoms. Fluorescence readings were taken using a SpectraMax i3x (Molecular Devices) with 485- and 535-mm excitation and emission wavelengths, respectively, at 30°C after 2.5 h of incubation.

### Tris-tricine-SDS Page

CFE reactions containing bacteriocins were mixed with Laemmli sample buffer (33 mM Tris-HCl, pH 6.8, 13 % glycerol, 2.1 % SDS, 0.01 % bromophenol blue) without DTT, directly loaded into a precast Tris-Tricine-SDS gel 16 % polyacrylamide (Invitrogen), run with Tris-MOPS-SDS running buffer (GenScript). After migration, gels were fixed in a solution of 10 % acetic acid and 20 % propanol, with agitation for 1h. The gel was washed with agitation, first with 70 % ethanol, a second time with 30 % ethanol and two last washes with Mili-Q water 1 hour each step. The washed gel was placed on top of dry LB solid (1.5 % agar) media in a square petri plate. A 600 µl sample from overnight cultured bacteria was mixed with LB solid media to complete 8 ml, then, poured on the LB solid plate with the gel and incubated overnight. Pictures were taken after incubation.

### Genome Sequencing

gDNA sequencing was performed with a MinION portable sequencing device (Oxford Nanopore). Briefly, gDNA samples were barcoded using a native barcoding kit 24 V14 (Oxford Nanopore), samples were loaded into a MinION flow cell (R10.4.1) following manufacturer’s instructions. Pod5 data were basecalled with dorado 0.7.3 with dna_r10.4.1_e8.2_400bps_sup@v5.0.0 model to generate fastq files. After basecalling, low quality reads (<Q12) were removed using Chopper [39]. and filtered reads were used as input to Breseq [40] with default parameters to identify genomic mutations.

### Toxicity and bacteriocin activity in Galleria mellonella

Adult larvae of *Galleria mellonella* (Terramania, Arnhem, Netherlands) were used for the infection model. Upon arrival, the larvae were stored at room temperature and used within three days. Healthy, non-discolored larvae weighing approximately 0.30 g were selected for analysis. The infected inoculum was prepared from an overnight pure culture on Columbia 5% sheep blood agar (Becton Dickinson, USA) and adjusted for each isolate to 10^6^ to 10^8^ CFU/mL to achieve 50% to 80% mortality within 48 hours. Ten larvae per group were inoculated by injecting a 10 µL aliquot into the hemocoel via the last left proleg using sterile 0.3 mL U100 insulin syringes (BD Micro-Fine) with a Microinjector (KDS 100 automated syringe pump, KD Scientific), followed by a 10 µL treatment or sterile saline serum injection (Mini-Plasco B.

Braun NaCl 0.9%) 15 minutes later on the right side. In each experiment, a group receiving two saline serum injections and another receiving the bacteriocin solution followed by a saline serum injection were used as negative and toxicity control groups, respectively. The insects were incubated in sterile Petri dishes, kept in the dark at 37°C in atmospheric air, and observed every 24 hours for 5 days. Health scores were calculated daily according to the health index score system [41], and larvae were considered dead if they did not respond to touch stimuli. Survival curves were plotted using the Kaplan-Meier method in GraphPad Prism 10. Statistical differences in survival rates were calculated using the Log-rank and Wilcoxon tests, with a significance level of *p* < 0.05.

## Acknowledgements

We thank Takashi Ebihara and Tomoko Miyagi from GeneFrontier Corporation for their assistance in the optimization of our DNA sequences. The work of AQ, KP and PG is founded by SWP Research from the Walloon Region (Grant N° 8723 PROACTIF). JB has financial support from contract (article 83) between Syngulon and the Universidad Complutense de Madrid (SYNGULON, S.A. / 27-2022) and Fondo Especial de Investigación (FEI) from the Universidad Complutense de Madrid (FEI24/08).

## Notes

### Competing Interest Statement

The authors have declared no competing interest.

## Bibliography

1. Cotter, P. D., Ross, R. P., & Hill, C. (2013). Bacteriocins—a viable alternative to antibiotics? Nature Reviews Microbiology, 11(2), 95–105.

2. Heilbronner, S., Krismer, B., Brötz-Oesterhelt, H., & Peschel, A. (2021). The microbiome-shaping roles of bacteriocins. Nature Reviews Microbiology, 19(11), 726–739.

3. Guinane, C. M., Lawton, E. M., O’Connor, P. M., O’Sullivan, Ó., Hill, C., Ross, R. P., & Cotter, P. D. (2016). The bacteriocin bactofencin A subtly modulates gut microbial populations. Anaerobe, 40, 41–49.

4. Rea, M.C., Dobson, A., O’Sullivan, O., Crispie, F., Fouhy, F., Cotter, P.D., Shanahan, F., Kiely, B., Hill, C. and Ross, R.P. (2011). Effect of broad-and narrow-spectrum antimicrobials on *Clostridium difficile* and microbial diversity in a model of the distal colon. Proceedings of the National Academy of Sciences, 108(supplement_1), 4639-4644.

5. Gebhart, D., Lok, S., Clare, S., Tomas, M., Stares, M., Scholl, D., Donskey, C.J., Lawley, T.D. and Govoni, G.R. (2015). A modified R-type bacteriocin specifically targeting *Clostridium difficile* prevents colonization of mice without affecting gut microbiota diversity. MBio, 6(2), 10–1128.

6. Gabant, P., & Borrero, J. (2019). PARAGEN 1.0: A standardized synthetic gene library for fast cell-free bacteriocin synthesis. Frontiers in Bioengineering and Biotechnology, 7, 213.

7. Peña, N., Bland, M.J., Sevillano, E., Muñoz-Atienza, E., Lafuente, I., Bakkoury, M.E., Cintas, L.M., Hernández, P.E., Gabant, P. and Borrero, J. (2022). *In vitro* and *in vivo* production and split-intein mediated ligation (SIML) of circular bacteriocins. Frontiers in Microbiology, 13, 1052686.

8. Jaumaux, F., Petit, K., Martin, A., Rodriguez-Villalobos, H., Vermeersch, M., Perez-Morga, D., & Gabant, P. (2023). Selective bacteriocins: a promising treatment for *Staphylococcus aureus* skin infections reveals insights into resistant mutants, vancomycin resistance, and cell wall alterations. Antibiotics, 12(6), 947.

9. Damoczi, J., Knoops, A., Martou, M.S., Jamaux, F., Gabant, P., Mahillon, J., Mignolet, J. and Hols, P. (2024). Uncovering the class II-bacteriocin predatiome in salivarius streptococci. bioRxiv, 2024-03

10. Pandi, A., Adam, D., Zare, A., Trinh, V. T., Schaefer, S. L., Burt, M., … & Erb, T. J. (2023). Cell-free biosynthesis combined with deep learning accelerates de novo-development of antimicrobial peptides. Nature Communications, 14(1), 7197.

11. Fedorec, A. J., Karkaria, B. D., Sulu, M., & Barnes, C. P. (2021). Single strain control of microbial consortia. Nature communications, 12(1), 1977.

12. Rutter, J.W., Dekker, L., Clare, C., Slendebroek, Z.F., Owen, K.A., McDonald, J.A., Nair, S.P., Fedorec, A.J. and Barnes, C.P. (2024). A bacteriocin expression platform for targeting pathogenic bacterial species. Nature Communications, 15(1), 6332.

13. Bartram, E., Asai, M., Gabant, P., & Wigneshweraraj, S. (2023). Enhancing the antibacterial function of probiotic *Escherichia coli* Nissle: when less is more. Applied and Environmental Microbiology, 89(11), e00975–23.

14. Coates, A. R., Hu, Y., Holt, J., & Yeh, P. (2020). Antibiotic combination therapy against resistant bacterial infections: synergy, rejuvenation and resistance reduction. Expert review of Anti-infective therapy, 18(1), 5–15.

15. Martin, A., Bland, M. J., Rodriguez-Villalobos, H., Gala, J. L., & Gabant, P. (2023). Promising antimicrobial activity and synergy of bacteriocins against *Mycobacterium tuberculosis*. Microbial Drug Resistance, 29(5), 165–174.

16. Mathur, H., Field, D., Rea, M. C., Cotter, P. D., Hill, C., & Ross, R. P. (2017). Bacteriocin-antimicrobial synergy: a medical and food perspective. Frontiers in microbiology, 8, 1205.

17. Rendueles, C., Duarte, A. C., Escobedo, S., Fernández, L., Rodríguez, A., García, P., & Martínez, B. (2022). Combined use of bacteriocins and bacteriophages as food biopreservatives. A review. International Journal of Food Microbiology, 368, 109611.

18. Vijayakumar, P. P., & Muriana, P. M. (2017). Inhibition of Listeria monocytogenes on ready-to-eat meats using bacteriocin mixtures based on mode-of-action. Foods, 6(3), 22.

19. Kranjec, C., Kristensen, S.S., Bartkiewicz, K.T., Brønner, M., Cavanagh, J.P., Srikantam, A., Mathiesen, G. and Diep, D.B. (2021). A bacteriocin-based treatment option for Staphylococcus haemolyticus biofilms. Scientific Reports, 11(1), 13909.

20. Guzmán, F., Barberis, S., & Illanes, A. (2007). Peptide synthesis: chemical or enzymatic. Electronic Journal of Biotechnology, 10(2), 279–314.

21. Garenne, D., Haines, M. C., Romantseva, E. F., Freemont, P., Strychalski, E. A., & Noireaux, V. (2021). Cell-free gene expression. Nature reviews methods primers, 1(1), 49.

22. Cintas, L. M., Casaus, P., Herranz, C., Håvarstein, L. S., Holo, H., Hernández, P. E., & Nes, I. F. (2000). Biochemical and genetic evidence that *Enterococcus faecium* L50 produces enterocins L50A and L50B, the sec-dependent enterocin P, and a novel bacteriocin secreted without an N-terminal extension termed enterocin Q. Journal of Bacteriology, 182(23), 6806–6814.

23. Izquierdo, E., Bednarczyk, A., Schaeffer, C., Cai, Y., Marchioni, E., Van Dorsselaer, A., & Ennahar, S. (2008). Production of enterocins L50A, L50B, and IT, a new enterocin, by *Enterococcus faecium* IT62, a strain isolated from Italian ryegrass in Japan. Antimicrobial agents and chemotherapy, 52(6), 1917–1923.

24. Schneider, T., Hahn-Löbmann, S., Stephan, A., Schulz, S., Giritch, A., Naumann, M., Kleinschmidt, M., Tusé, D. and Gleba, Y. (2018). Plant-made *Salmonella* bacteriocins salmocins for control of *Salmonella* pathovars. Sci Rep 8: 4078.

25. Telhig, S., Ben Said, L., Zirah, S., Fliss, I., & Rebuffat, S. (2020). Bacteriocins to thwart bacterial resistance in gram negative bacteria. Frontiers in microbiology, 11, 586433.

26. HÃ¥ varstein, L. S., Holo, H., & Nes, I. F. (1994). The leader peptide of colicin V shares consensus sequences with leader peptides that are common among peptide bacteriocins produced by Gram-positive bacteria. Microbiology, 140(9), 2383–2389.

27. Drusano, G.L., Neely, M., Van Guilder, M., Schumitzky, A., Brown, D., Fikes, S., Peloquin, C. and Louie, A. (2014). Analysis of combination drug therapy to develop regimens with shortened duration of treatment for tuberculosis. PloS one, 9(7), e101311.

28. Zhou, A., Kang, T.M., Yuan, J., Beppler, C., Nguyen, C., Mao, Z., Nguyen, M.Q., Yeh, P. and Miller, J.H. (2015). Synergistic interactions of vancomycin with different antibiotics against Escherichia coli: trimethoprim and nitrofurantoin display strong synergies with vancomycin against wild-type *E. coli*. Antimicrobial agents and chemotherapy, 59(1), 276–281.

29. Coates, A. R., Hu, Y., Holt, J., & Yeh, P. (2020). Antibiotic combination therapy against resistant bacterial infections: synergy, rejuvenation and resistance reduction. Expert review of Anti-infective therapy, 18(1), 5–15.

30. Noireaux, V., Bar-Ziv, R., & Libchaber, A. (2003). Principles of cell-free genetic circuit assembly. Proceedings of the National Academy of Sciences, 100(22), 12672–12677.

31. Cappuccio, J.A., Blanchette, C.D., Sulchek, T.A., Arroyo, E.S., Kralj, J.M., Hinz, A.K., Kuhn, E.A., Chromy, B.A., Segelke, B.W., Rothschild, K.J. and Fletcher, J.E. (2008). Cell-free co-expression of functional membrane proteins and apolipoprotein, forming soluble nanolipoprotein particles. Molecular & Cellular Proteomics, 7(11), 2246–2253.

32. Terada, T., Murata, T., Shirouzu, M., & Yokoyama, S. (2014). Cell-free expression of protein complexes for structural biology. Structural Genomics: General Applications, 151-159.

33. Numata, K., Motoda, Y., Watanabe, S., Osanai, T., & Kigawa, T. (2015). Co-expression of two polyhydroxyalkanoate synthase subunits from *Synechocystis* sp. PCC 6803 by cell-free synthesis and their specific activity for polymerization of 3-hydroxybutyryl-coenzyme A. Biochemistry, 54(6), 1401–1407.

34. Park, Y. J., Lee, K. H., & Kim, D. M. (2017). Assessing translational efficiency by a reporter protein co-expressed in a cell-free synthesis system. Analytical biochemistry, 518, 139–142.

35. Liu, W.Q., Ji, X., Ba, F., Zhang, Y., Xu, H., Huang, S., Zheng, X., Liu, Y., Ling, S., Jewett, M.C. and Li, J. (2024). Cell-free biosynthesis and engineering of ribosomally synthesized lanthipeptides. Nature Communications, 15(1), 4336.

36. Emslander, Q., Vogele, K., Braun, P., Stender, J., Willy, C., Joppich, M., … & Westmeyer, G. G. (2022). Cell-free production of personalized therapeutic phages targeting multidrug-resistant bacteria. Cell chemical biology, 29(9), 1434–1445.

37. Sword, T. T., Abbas, G. S., & Bailey, C. B. (2024). Cell-free protein synthesis for nonribosomal peptide synthetic biology. Frontiers in Natural Products, 3, 1353362.

38. Baba, T., Ara, T., Hasegawa, M., Takai, Y., Okumura, Y., Baba, M., Datsenko, K.A., Tomita, M., Wanner, B.L. and Mori, H. (2006). Construction of *Escherichia coli K-12* in-frame, single-gene knockout mutants: the Keio collection. Molecular systems biology, 2(1), 2006–0008.

39. De Coster, W., & Rademakers, R. (2023). NanoPack2: population-scale evaluation of long-read sequencing data. Bioinformatics, 39(5), btad311.

40. Deatherage, D. E., & Barrick, J. E. (2014). Identification of mutations in laboratory-evolved microbes from next-generation sequencing data using breseq. Engineering and analyzing multicellular systems: methods and protocols, 165–188.

41. Ménard, G., Rouillon, A., Cattoir, V., & Donnio, P. Y. (2021). *Galleria mellonella* as a suitable model of bacterial infection: past, present and future. Frontiers in cellular and infection microbiology, 11, 782733.

