## Supplementary data for "Multiplexing bacteriocin synthesis to kill and prevent antimicrobial resistance"

DNA sequences before and after optimization with template DNA guidelines from PureFrex (GeneFrontier Corporation)

T7 promoter

Lac operator

RBS

Gene

T7 terminator

rrnB terminator

Original gene device for GFP expression

TAATACGACTCACTATAGGGGAATTGTGAGCGGATAACAATTCCCCTCTAGAAATAATTTTGTTTAACTTTAAGAAG
GAGATATACATATGCGTAAAGGTGAAGAGTTGTTTACAGGTGTAGTGCCCATCCTGGTCGAGCTTGATGGCGACG
TGAATGGTCATAAGTTCTCAGTTAGCGGTGAAGGTGAGGGCGATGCGACGTATGGGAAACTCACCCTCAAATTTA
TTTGTACAACTGGCAAATTACCGGTCCCGTGGCCCACGTTAGTTACCACATTCGGTTATGGCGTTCAGTGCTTCGC
ACGTTATCCGGACCACATGAAACGTCATGACTTCTTCAAATCCGCGATGCCTGAAGGTTACGTCCAAGAACGTACT
ATCTTTTTTAAAGACGATGGCAACTATAAAACCCGGGCCGAGGTTAAGTTTGAAGGCGATACCCTGGTGAATCGT
ATCGAACTGAAAGGTATTGACTTTAAAGAAGATGGCAACATACTGGGCCATAAGCTGGAATACAACTATAACAGC
CATAACGTCTATATCATGGCGGATAAGCAGAAAAACGGTATTAAAGTTAATTTCAAGATTCGACACAATATTGAAG
ACGGTTCGGTACAACTGGCCGATCATTACCAGCAGAACACACCGATCGGCGACGGTCCTGTTTTACTGCCCGATA
ATCACTATCTGTCAACTCAGTCGGCGTTGTCTAAGGACCCGAATGAAAAACGCGATCACATGGTATTGCTTGAATT
TGTGACTGCCGCGGGTATCACGCACGGTATGGACGAACTTTATAAGTAGGATCCATCCGGCTGCTAACAAAGCCC
GAAAGGAAGCTGAGTTGGCTGCTGCCACCGCTGAGCAATAACTAGCATAACCCCTTGGGGCCTCTAAACGGGTC
TTGAGGGGTTTTTTGCTGAAAGGAGGAACTATATCCGGATTCAGGAGAGCGTTCACCGACAAACAACAGATAAA
ACGAAAGGCCCAGTCTTTCGACTGAGCCTTTCGTTTTATTTG

Optimized gene device for GFP expression.

GAAATTAATACGACTCACTATAGGGAGACCACAACGGTTTCCCTCTAGAAATAATTTTGTTTAACTTTAAGAAGGA
GATATACCAATGCGAAAAGGAGAAGAATTATTTACAGGTGTGGTTCCTATCCTCGTAGAGCTGGACGGTGACGTT
AACGGCCATAAGTTCTCTGTCTCCGGTGAAGGCGAAGGTGATGCTACCTATGGGAAACTTACTCTGAAATTTATTT
GTACCACAGGCAAACTGCCGGTGCCCTGGCCGACTCTGGTTACGACCTTCGGTTACGGCGTACAGTGCTTCGCG
CGCTACCCGGACCACATGAAGCGTCACGACTTCTTCAAAAGTGCTATGCCAGAAGGTTATGTTCAAGAGCGCACT
ATCTTCTTTAAAGACGATGGAAACTACAAAACCCGTGCAGAAGTGAAGTTCGAAGGTGACACTTTGGTAAATCGT
ATCGAGCTGAAAGGCATTGATTTCAAAGAAGACGGTAACATCTTAGGGCATAAACTGGAATACAACTATAACAGC
CACAATGTTTACATCATGGCCGATAAGCAGAAAAACGGCATTAAAGTCAACTTTAAAATCCGCCATAACATCGAAG
ACGGTTCTGTTCAGCTCGCTGATCACTACCAACAGAATACGCCTATTGGCGACGGTCCGGTGCTGCTTCCGGATA
ACCACTATCTGTCCACCCAGAGCGCGCTGTCTAAGGACCCGAACGAGAAACGTGATCATATGGTACTGCTGGAAT
TCGTTACTGCAGCTGGCATCACCCACGGTATGGACGAACTGTACAAATAAGATCCATCCGGCTGCTAACAAAGCC
CGAAAGGAAGCTGAGTTGGCTGCTGCCACCGCTGAGCAATAACTAGCATAACCCCTTGGGGCCTCTAAACGGGT
CTTGAGGGGTTTTTTGCTGAAAGGAGGAACTATATCCGGATTCAGGAGAGCGTTCACCGACAAACAACAGATAA
AACGAAAGGCCCAGTCTTTCGACTGAGCCTTTCGTTTTATTTG

Original gene device for MccV expression

TAATACGACTCACTATAGGGGAATTGTGAGCGGATAACAATTCCCCTCTAGAAATAATTTTGTTTAACTTTAAGAAG
GAGATATACATATGGCGTCCGGCCGCGACATCGCGATGGCGATCGGTACTTTAAGCGGGCAATTTGTCGCAGGCG
GTATCGGCGCGGCCGCGGGGGGCGTTGCCGGCGGTGCTATTTATGATTATGCATCAACCCATAAACCGAATCCCG
CCATGAGTCCATCGGGACTGGGTGGCACGATTAAACAGAAGCCTGAAGGTATTCCGTCGGAGGCCTGGAACTAC
GCAGCTGGACGTTTGTGTAATTGGTCTCCGAACAACCTCAGCGATGTGTGCCTGTAGGATCCATCCGGCTGCTAA
CAAAGCCCGAAAGGAAGCTGAGTTGGCTGCTGCCACCGCTGAGCAATAACTAGCATAACCCCTTGGGGCCTCTA
AACGGGTCTTGAGGGGTTTTTTGCTGAAAGGAGGAACTATATCCGGATTCAGGAGAGCGTTCACCGACAAACA
ACAGATAAAACGAAAGGCCCAGTCTTTCGACTGAGCCTTTCGTTTTATTTG

Optimized gene device for MccV expression

GAAATTAATACGACTCACTATAGGGAGACCACAACGGTTTCCCTCTAGAAATAATTTTGTTTAACTTTAAGAAGGA
GATATACCAATGGCTTCAGGTCGAGATATTGCAATGGCAATTGGTACCCTGTCCGGCCAGTTCGTGGCGGGTGGA
ATCGGTGCAGCTGCGGGCGGTGTAGCTGGGGGCGCAATCTACGATTATGCCAGTACTCACAAGCCGAACCCAGC
TATGAGCCCTTCTGGTCTGGGCGGTACCATTAAACAGAAACCGGAAGGCATCCCGTCCGAAGCGTGGAACTACG
CAGCTGGTCGTCTGTGTAATTGGAGCCCGAACAACCTGTCTGACGTTTGCCTGTAGGATCCATCCGGCTGCTAAC
AAAGCCCGAAAGGAAGCTGAGTTGGCTGCTGCCACCGCTGAGCAATAACTAGCATAACCCCTTGGGGCCTCTAA
ACGGGTCTTGAGGGGTTTTTTGCTGAAAGGAGGAACTATATCCGGATTCAGGAGAGCGTTCACCGACAAACAAC
AGATAAAACGAAAGGCCCAGTCTTTCGACTGAGCCTTTCGTTTTATTTG
